# Cyanobacterial metagenomes reflect the spatiotemporal variations in a coastal brackish lagoon

**DOI:** 10.1101/2025.04.10.648224

**Authors:** Manisha Ray, Govindhaswamy Umapathy

## Abstract

Cyanobacteria play vital roles in aquatic ecosystems by driving photosynthesis, nitrogen fixation, carbon sequestration, and forming symbiotic relationships with diverse organisms. However, their proliferation can trigger harmful algal blooms, posing risks to aquatic biodiversity and public health. Despite their ecological significance, the interplay between cyanobacterial genomic traits and ecosystem dynamics remains poorly resolved. Here, we employed culture-independent metagenomic approaches to reconstruct cyanobacterial metagenome-assembled genomes (MAGs) from Chilika Lagoon, India, and investigate their spatiotemporal distribution and functions. Our analysis revealed distinct temporal patterns in cyanobacterial MAG abundance, with salinity emerging as the primary environmental driver of community structure and functional gene composition. Genes associated with biogeochemical cycling and toxin synthesis displayed pronounced seasonal variation, suggesting that functional genomic traits, rather than taxonomic identity govern species selection. Notably, five MAGs harboured the complete phosphate acetyltransferase-acetate kinase (Pta-Ack) pathway, a critical component of the Wood–Ljungdahl pathway, indicating an underappreciated potential for alternative carbon fixation mechanisms alongside the canonical Calvin-Benson-Bassham cycle. Furthermore, genomic variability, rather than phylogenetic relatedness was the dominant factor shaping cyanobacterial dynamics in the lagoon. This study establishes a direct link between physicochemical fluctuations and cyanobacterial functional diversity, offering critical insights into how climate-driven changes in salinity and nutrient regimes may influence aquatic ecosystems. By elucidating the genomic basis of cyanobacterial adaptation, these findings enhance our capacity to predict ecological outcomes of harmful algal blooms and inform strategies to safeguard ecosystem services in vulnerable coastal habitats.

**Importance:** This study employs culture-independent metagenomics to reconstruct cyanobacterial metagenome-assembled genomes (MAGs) in Chilika Lagoon, India, unraveling their spatiotemporal distribution and functional traits. Salinity emerged as the primary driver shaping community structure and functional gene composition, with seasonal fluctuations influencing genes tied to biogeochemical cycling (e.g., carbon, nitrogen) and toxin synthesis. Notably, functional genomic traits—rather than taxonomic identity—governed species selection, highlighting adaptive strategies under environmental stress. Intriguingly, five MAGs harbored the complete Pta-Ack pathway, a component of the Wood-Ljungdahl pathway, suggesting cyanobacteria may employ alternative carbon fixation mechanisms alongside the Calvin-Benson-Bassham cycle under fluctuating conditions. These findings link physicochemical variables (e.g., salinity, nutrients) to functional diversity, revealing how genomic adaptations underpin ecological resilience. The study provides critical insights into cyanobacterial responses to environmental change by bridging microbial genomic plasticity with ecosystem-level impacts. This framework aids in predicting bloom dynamics and toxin risks, offering actionable tools to mitigate ecological threats in vulnerable coastal habitats, thereby informing conservation and management strategies amid climate variability.

## 1. Introduction

Cyanobacteria are the only prokaryotes that can perform oxygenic photosynthesis (Sánchez-Baracaldo et al., 2022). They account for 20–30% of Earth’s photosynthetic productivity as primary producers (Waterbury et al., 1979). Cyanobacteria play critical roles in supporting ecosystem services such as biogeochemical cycling by acting as biological drivers of nitrogen and phosphorus cycling in lakes (Ottingham et al., 2015). They constitute the key group among bacteria that utilize the Calvin– Benson–Bassham pathway for carbon fixation (Berg, 2011).

The variability of cyanobacterial species with environmental heterogeneity is only partially understood. A few studies have identified anthropogenic disturbances in the form of eutrophication of aquatic bodies as driving factors that favour certain cyanobacterial species. These are associated mainly with bloom-forming cyanobacterial genera such as *Microcystis, Cylindrospermopsis and Dolichospermum* (Dalu and Wasserman, 2018; Tromas et al., 2017) These studies have critical implications for understanding the impact of climate change on aquatic bodies (O’Neil et al., 2012). Studies have also focused on lake stratification with seasonal changes and their impact on cyanobacterial community composition, which is driven by nutrients and light (Nwosu et al., 2021). Furthermore, certain cyanobacterial species act as indicators of environmental conditions (due to an increase in certain micronutrient levels in the water body) (Cottingham et al., 2015). They can thus assist in tracking the health of these aquatic ecosystems.

In most of these cases, a conserved and hypervariable region of the genome is chosen to elucidate the bacterial diversity of the ecosystem and correlate it with other varying abiotic parameters. For example, (Kraemer et al., 2020) assessed the ecological impact of land use on the bacterial diversity, community and composition of several lakes in Eastern Canada. Only a handful of studies from lakes in India have evaluated bacterial diversity with respect to spatiotemporal heterogeneity and abiotic characteristics. One such interesting aquatic ecosystem is a biodiverse and largest brackish coastal lagoon in Asia, the Chilika lagoon (Dash et al., 2024) (Mohapatra et al., 2020). Such studies are crucial for understanding the impacts of proximate causes such as biogeography, spatiotemporal shifts in abiotic parameters, anthropogenic disturbances and climate change on the ecosystem, which favour the proliferation of one bacterial species over another.

Although there are substantial studies delineating the abiotic drivers of cyanobacterial heterogeneity in space and time, there is a paucity of understanding of the ultimate reasons behind these patterns. The bacterial species that can thrive and increase in environmental conditions depend on the genes present in the organism. Hence, the organism’s genome serves as a toolkit that aids bacterial species in adapting and multiplying at a particular physiochemical condition of the ecosystem. In certain studies, the genomic component is inferred from bacterial diversity by predicting functional metabolic pathways of the closest related species with genomes available in public databases. In a study by (Ramoneda et al., 2021), the connectivity between rivers and their habitat conditions was studied to decipher the changes in microbial assemblages. The functional gene potential of these genes was extrapolated from their 16S rRNA sequences. With the rise of high-throughput sequencing and advancements in computational resources, finding the ultimate cause behind certain species being favoured over others seems more achievable via culture-independent methods of bacterial genome assembly. Hence, deciphering the bacterial community composition can be beneficial in two ways: identifying the broad physiochemical conditions of water bodies and identifying the genes required to thrive under those conditions. This bridges the spatiotemporal changes in the physiochemical conditions of the ecosystem and the genomic traits of bacteria necessary to flourish there.

Our study was conducted in Chilika Lagoon, which is located on the eastern coast of the Indian subcontinent and is a biodiverse yet understudied region. In this study, we aimed to identify the major abiotic drivers of cyanobacterial species and the functions encoded by their metagenomes in the brackish coastal lagoon of Chilika. We further aimed to relate the genomic traits of cyanobacteria, particularly those involved in biogeochemical cycling and secondary metabolite production, with the impact of seasonal variations over the spatial scale of Chilika Lagoon. We also aimed to explore whether the variations in cyanobacterial composition were linked to their phylogenetic relatedness.

## 2. Materials and methods

### 2.1 Study site, eDNA isolation and assembly of cyanobacterial metagenomes

Chilika Lagoon (19° 28′ N: 19° 54′ N and 85° 06′ E: 85°35′ E) is the largest brackish water lagoon in Asia, and because of its high biodiversity, the first wetland was protected under the Ramsar Convention of 1971. This lagoon is located on the eastern coast of India and is connected to the Bay of Bengal. The hydrology of the lagoon varies with season and between years (Finlayson et al., 2020), which also leads to changes in the physicochemical parameters of the lagoon. The hydrology of the lagoon also dictates the inflow of inorganic N and P nutrients, promoting primary productivity in lagoon regions (Mukherjee et al., 2019). The physiochemical parameters (water temperature, pH, transparency, depth, biological oxygen demand (BOD), dissolved oxygen (DO), phosphate, silicate and nitrate) were measured for the water samples from all the sampling sites described in Ray et al. 2024 (Ray et al., 2024).

At the water sampling sites of Chilika Lagoon, extraction and sequencing of eDNA were performed as previously described by Manu and Umapathy 2023 (Manu and Umapathy, 2023). In brief, 9 unique locations (16 samples in triplicates) were filtered with a minimum 10 litres of water. eDNA was isolated from these mixed-cellulose ester membranes and was prepared for sequencing via the Illumina TruSeq DNA PCR-free library preparation method and sequenced over 300 cycles on the NovaSeq 6000 platform. The assembly of reads and taxonomic annotation was carried out as described in Ray et al. 2024 (Ray et al., 2024). In summary, the obtained raw reads were trimmed, filtered on the basis of quality, normalized, error-corrected and coassembled from all samples. The contigs that were more than 1000 bp in length were binned and the relative abundance of each MAG in every season and sampling site was calculated.

### 2.2 Cyanobacterial MAGs and their orthogroup search

Three sampling seasons (winter, summer and monsoon) had several sampling sites. Winter sampling was performed for two years, 2019 and 2020. This occurred because the winter 2019 sampling was performed as a pilot study, which was followed by sampling the following year in three different seasons. On the basis of the presence and absence of cyanobacterial MAGs in this season, a Venn diagram was constructed using Venny 2.1(Oliveros JC, 2007-2015).

We predicted all the protein-coding genes from the cyanobacterial MAGs using Prodigal.v2.6.3 (Hyatt et al., 2010) with the meta option. Each MAG was searched for orthogroups (genes that originated from a single ancestral gene in the last common ancestor of a species clade) using Orthofinder version 2.5.4 (Emms and Kelly, 2019). We plotted the number of orthogroups in each MAG using the output from Orthofinder and executed it in RStudio with the ggplot2 v.3.5.0 (Wickham, 2016) and readr v.2.1.5 (Wickham H, Hester J, 2024)s packages. We also generated a plot for the shared orthogroups in MAG pairs using heatmap.2 function from gplots version 3.1.3.1 (Warnes et al., 2024). We further converted the matrix for the orthogroups present in a MAG to a binary matrix based on the presence and absence of orthogroups in each MAG to account for biases due to the differential incompleteness of MAGs.

### 2.3 Phylogenetic analysis

We used all 14 high-quality MAGs (photosynthetic) and two non-photosynthetic cyanobacterial MAGs to extract single-copy orthologues using Orthofinder version 2.5.4. This occurred because the medium-quality MAGs were partially complete and hence some orthogroups were missing, so they were not included in the phylogenetic construction. The single-copy orthogroups were individually aligned using MAFFT v7.505 (Katoh and Standley, 2013), trimmed using trimal v1.4.rev15 (Capella-gutiérrez et al., 2009) and concatenated using amas (Borowiec, 2016). The partitions and concatenated files were then used as inputs for IQ-TREE multicore version 2.3.2 (Minh et al., 2020) for maximum likelihood tree construction. The tree was visualized using iTOL v6 (Letunic and Bork, 2024). A heatmap for the relative abundances of cyanobacterial MAGs in various seasons was constructed using RStudio and the ggplot2 (Wickham, 2016) and reshape2 (Wickham, 2007) packages.

### 2.4 Statistical analysis

We used redundancy analysis (RDA) to correlate changes in cyanobacterial composition with gradients in the abiotic environment. First, we checked for pairwise correlations among the various abiotic parameters using Pearson correlation (corrplot package v.0.92 (Simko, 2021)). Variables with correlation coefficients ≥ 0.5 were not considered together in the RDA model. We did not include the summer season in this analysis, as logistical issues precluded the collection of data for certain abiotic variables. The winters of 2019 and 2020 were considered one group.

#### 2.4.1 Relative abundances of MAGs as response variables and physiochemical factors as explanatory variables

We performed the Hellinger transformation using BiodiversityR version 2.15-4 (Kindt R, 2005) of the relative abundances of MAGs to normalize the data by standardizing the abundances to MAG totals and then square-rooting them. The rda function from the vegan v.2.6-4 package (Oksanen et al., 2022) in RStudio was used to compute the relative abundance of cyanobacterial MAGs as the response variable, and physiochemical parameters (selected based on corrplot) were used as the explanatory variable. We used the function ordiR2step to build the forward model to maximize the adjusted R^2^ value and the ordiplot function from the vegan package v.2.6-4 to plot the RDA results. We used the ggordiplots package v.0.4.3 (Quensen J Simpson G, 2024) to create an ordination plot with group ellipses based on season, with a confidence of 0.95.

#### 2.4.2 Presence-absence of Cluster of Orthologous Groups (COGs) as response variables and physiochemical factors as explanatory variables

We used a custom Python script to compute the COG-site matrix using binary matrices of COG-MAG and MAG sites as the inputs. The COG-site matrix was further converted to a binary matrix to address the biases associated with the quality of draft MAGs. The function rda() from vegan v.2.6-4 was subsequently used with the presence-absence of COGs at sites as the response variable and physiochemical parameters (Salinity+BOD+Transparency+Phosphate+Silicate) as explanatory variables. Furthermore, the function ordiR2step was used to build a forward model such that it maximizes the adjusted R^2^ value. The ordiplot function from the vegan package v.2.6-4 was used to visualize the RDA results. The ggordiplots package v.0.4.3 was used to make an ordination plot with group ellipses based on season, with a confidence of 0.95.

#### 2.4.3 Relationships between cyanobacterial MAGs and functional genes relevant to biogeochemical cycling

The predicted ORFs were annotated based on their KEGG IDs, which represent functional traits. The MAGs present in a minimum of 50% of all the sampled sites of a particular season were grouped. The KEGG IDs for biogeochemically important functions (nitrogen metabolism, phosphate metabolism, sulfur metabolism, carbon fixation and photosynthesis) were searched in all these groups of MAGs. The mean number of ORFs for each functional trait was calculated (the KEGG pathway was used as a reference for the keg IDs of each pathway). We used ggplot2 (Wickham, 2016) to visualize the results and to draw inferences.

For nitrogen metabolism, cyanobacterial MAGs with ORFs for genes involved in nitrogen fixation, dissimilatory nitrate reduction and assimilatory nitrate reduction were considered. For phosphate metabolism, MAGs with ORFs for genes responsible for P-starvation response regulation, P-uptake and transport systems and inorganic P-solubilization and organic P-mineralization were considered. Carbon fixation in prokaryotes was associated with seven pathways in KEGG, and the MAGs with ORFs for the reductive citrate cycle, dicarboxylate-hydroxybutyrate cycle, hydroxypropionate-hydroxybutylate cycle, 3-hydroxypropionate bicycle, reductive acetyl-CoA pathway (Wood‒ Ljungdahl pathway), phosphate acetyltransferase-acetate kinase pathway and incomplete reductive citrate cycle (acetyl-CoA => oxoglutarate) were considered for analysis. Photosynthesis has four major components in its pathway, which were considered for further analysis. These genes were F-type ATPases, photosystem II, the cytochrome b6f complex and photosystem I. We calculated Friedman rank sum test using baseR and function friedman.test() to check for statistical significance of various pathways across seasons.**2.4.4 Secondary metabolite prediction in cyanobacterial MAGs and their seasonal variations**

We used antiSMASH v.6.0 (Blin et al., 2021) to identify and annotate gene clusters involved in secondary metabolite biosynthesis. The results were then processed with BiG-SCAPE v.1.1.8 (https://bigscape-corason.secondarymetabolites.org/) applying a gene cluster family (GCF) clustering threshold of 0.3 and using the MIBiG reference database version 2.1 (Medema et al., 2015) to cluster the sequences into systematic BGC categories. ggplot2 (Wickham, 2016) was used to plot the distribution of BGC across seasons. To test the statistical significance of various BGC category counts across seasons, a chi-square test was performed in RStudio using the function chisq.test from the package stats version v4.2.2 (Team, 2022).

## 3. Results

### 3.1 Spatiotemporal distribution of cyanobacterial MAGs and their phylogenetic relatedness

We assembled a total of 83 cyanobacterial MAGs which belonged to 15 different families (9 MAGs were unclassified at family-level). Out of these 83 MAGs, 14 were high-quality and the remaining were medium-quality, on the basis of MIMAG guidelines ((Bowers et al., 2017)Bowers et al., 2017). Interestingly, 22 cyanobacterial MAGs (Phormidesmiacea, Microcystaceae_A, Cyanobiaceae and a family of non-photosynthetic Sericytochromatia) were present in all four sampling seasons. Most of them were from familiy Cyanobiaceae (18), of which 7 were from genus Cyanobium. The 2 MAGs from family Phormidesmiaceae belonged to genus Nodosilinea and a MAG from family Microcystaceae was from the Synechocystis genera.

A total of 30 MAGs were present in only a specific season. 27 cyanobacterial MAGs [Cyanobiaceae (10), Nostocaceae (4), Microcystaceae_A (3), Phormidesmiaceae (2), Elainellaceae (1), Leptolyngbyaceae (1), Microcoleaceae (1), Prochlorotrichaceae (1)] were exclusively found in the winter of 2019, whereas there were no new MAGs in the following winter season. Also, there were 2 unique MAGs in monsoon of 2020 and 1 unique MAG in summer of 2020 (Figure 1).

**Figure 1:**
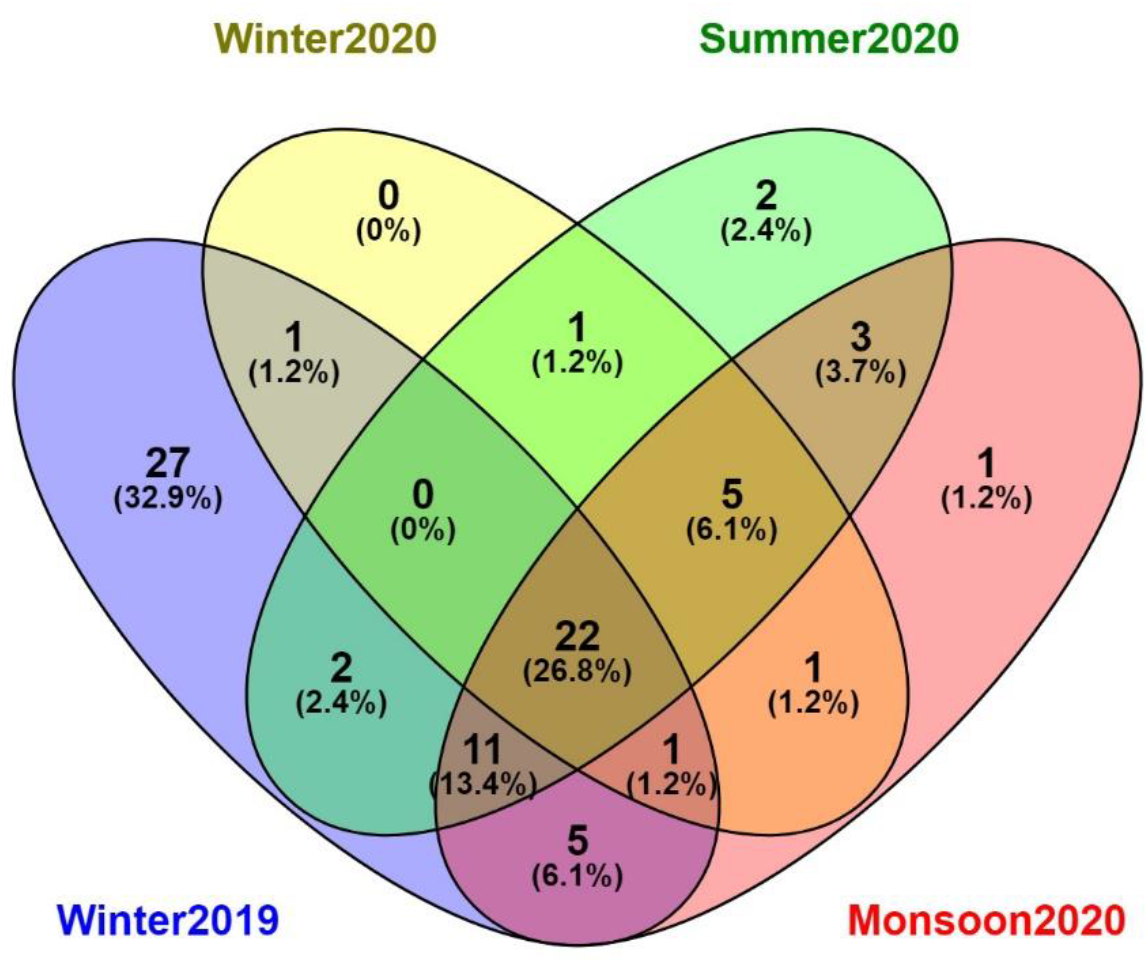
Venn diagram of the seasonal distribution of cyanobacterial MAGs

Pearson correlations were calculated between the physiochemical parameters of Chilika Lagoon (Supplementary 3) which showed that the model selected was ‘Salinity + BOD + Transparency + Phosphate + Silicate’. The RDA revealed that environmental variables accounted for 55.93% of the variance in the relative abundance of cyanobacterial MAGs (Figure 2).

**Figure 2:**
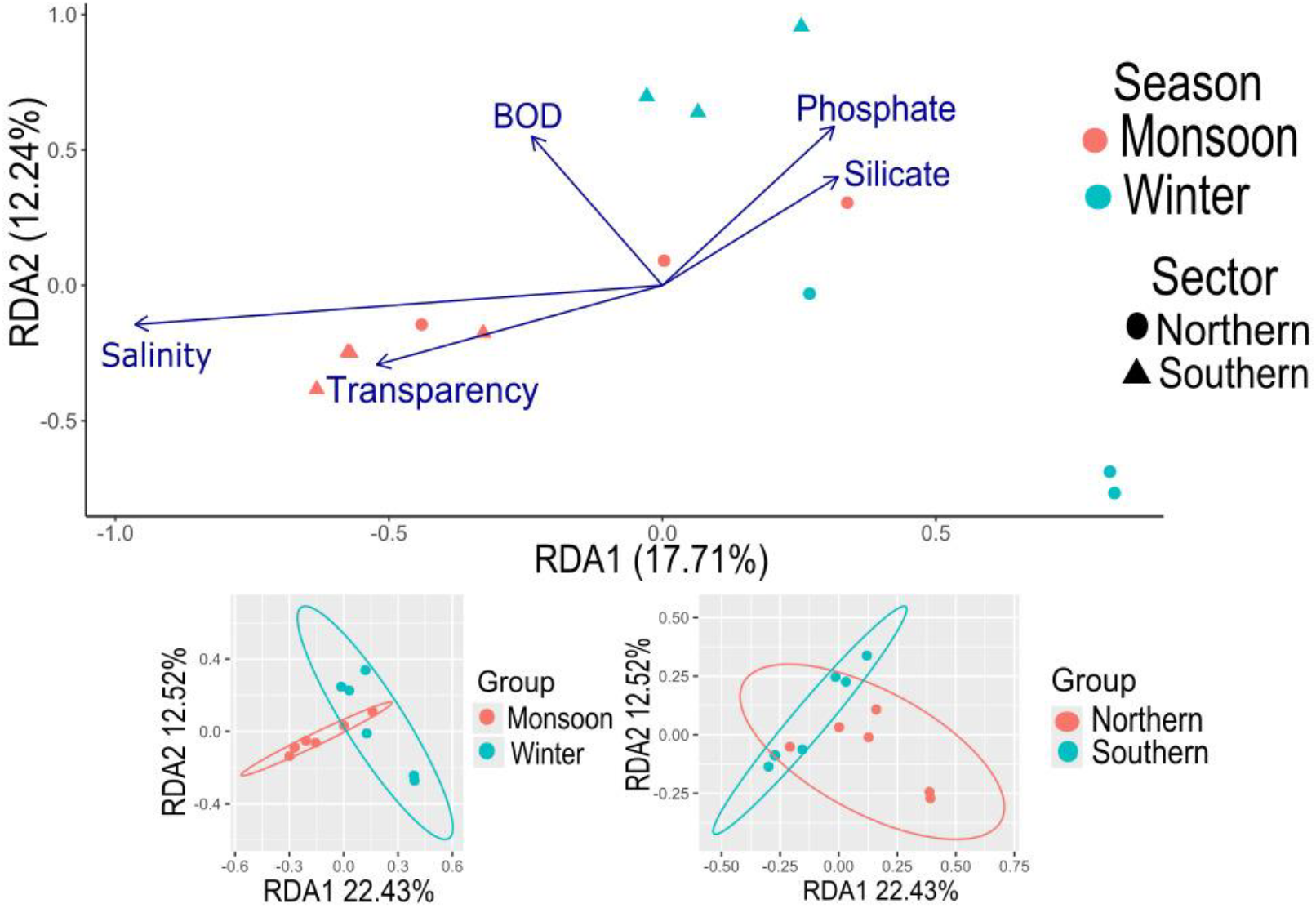
RDA with physicochemical parameters as explanatory variables and the relative abundances of cyanobacterial MAGs as response variables

After running OrthoFinder on 16 cyanobacterial MAGs from Chilika Lagoon, we identified 130 single-copy orthologues. The phylogenetic tree generated using these orthologues exhibited a topology consistent with their GTDB-assigned taxonomy. However, the relative abundances of MAGs within the same family showed little concordance in most cases (Figure 3).

**Figure 3:**
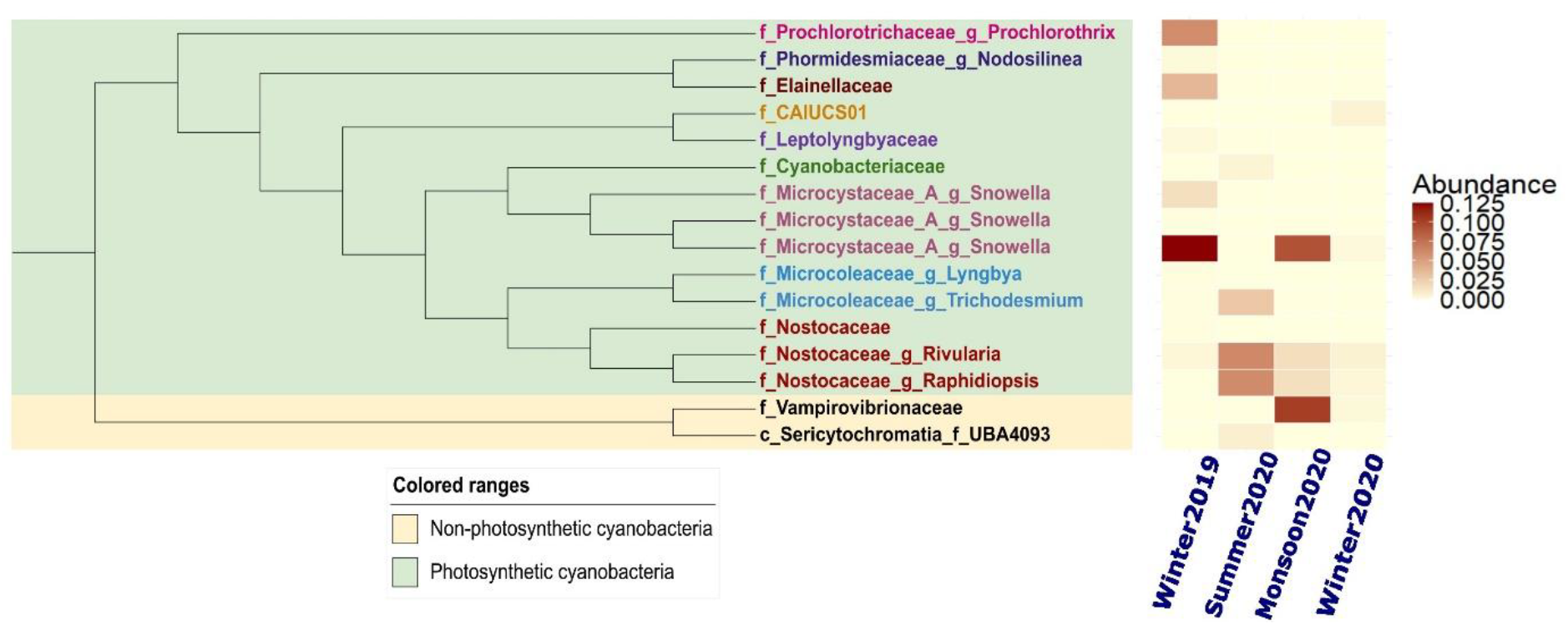
Maximum likelihood tree constructed using single-copy orthologues from cyanobacterial MAGs and their relative abundances across the sampled seasons

### 3.2 Spatiotemporal distribution of orthogroups

A total of 11,822 orthogroups were predicted from the 83 MAGs, of which 757 were species-specific. Nine orthogroups were present in all MAGs and can be considered core orthogroups. The number of orthogroups was highest for the high-quality MAGs, except for some which were medium-quality drafts (Supplementary 1). This reflects that the quality of the draft is not the sole criterion for the number of COGs present in the MAGs. The number of COGs was also influenced by the taxonomic level, with certain taxa having more COGs than others (Supplementary 2).

Redundancy analysis revealed that salinity explained most of the variation in the relative abundances of cyanobacterial MAGs, with a statistical significance of 0.002. The RDA results indicated that the environmental variables explained 45.34% of the variance in the presence and absence of COGs in the cyanobacterial MAGs (Figure 4). Salinity explained most of the variance in the presence and absence of COGs in cyanobacterial MAGs, among other abiotic variables. We also found that the space occupied by the ellipses differed by season.

**Figure 4:**
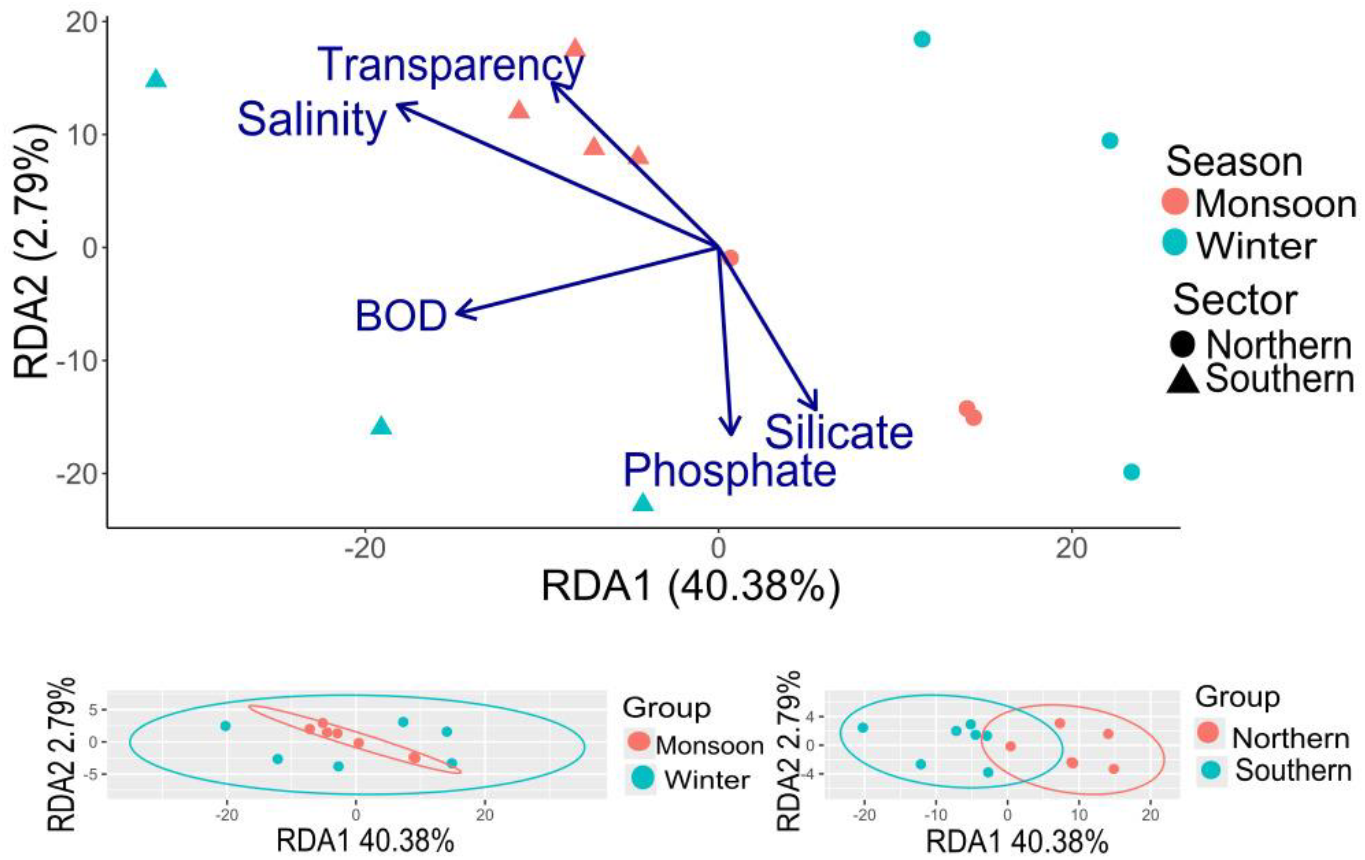
RDA with physicochemical parameters as explanatory variables and the presence-absence of COGs in cyanobacterial MAGs as response variables

### 3.3 Temporal shifts in nitrogen metabolism, phosphate metabolism, photosynthesis and carbon fixation pathways in the cyanobacterial MAGs

The various nitrogen metabolism pathways varied temporally (Friedman test, p-value = 0.0387) and the highest mean ORFs were found in the winter of 2019 (Figure 5). Cyanobacterial species present at the sampling sites in the winter of 2019 presented markedly greater mean ORFs of nitrogen metabolism pathways (Figure 6). Further, dissimilatory and assimilatory nitrate reduction exhibited opposing trends across seasons. Monsoon had the highest mean ORFs for assimilatory nitrate reduction but the lowest for dissimilatory nitrate reduction.

**Figure 5:**
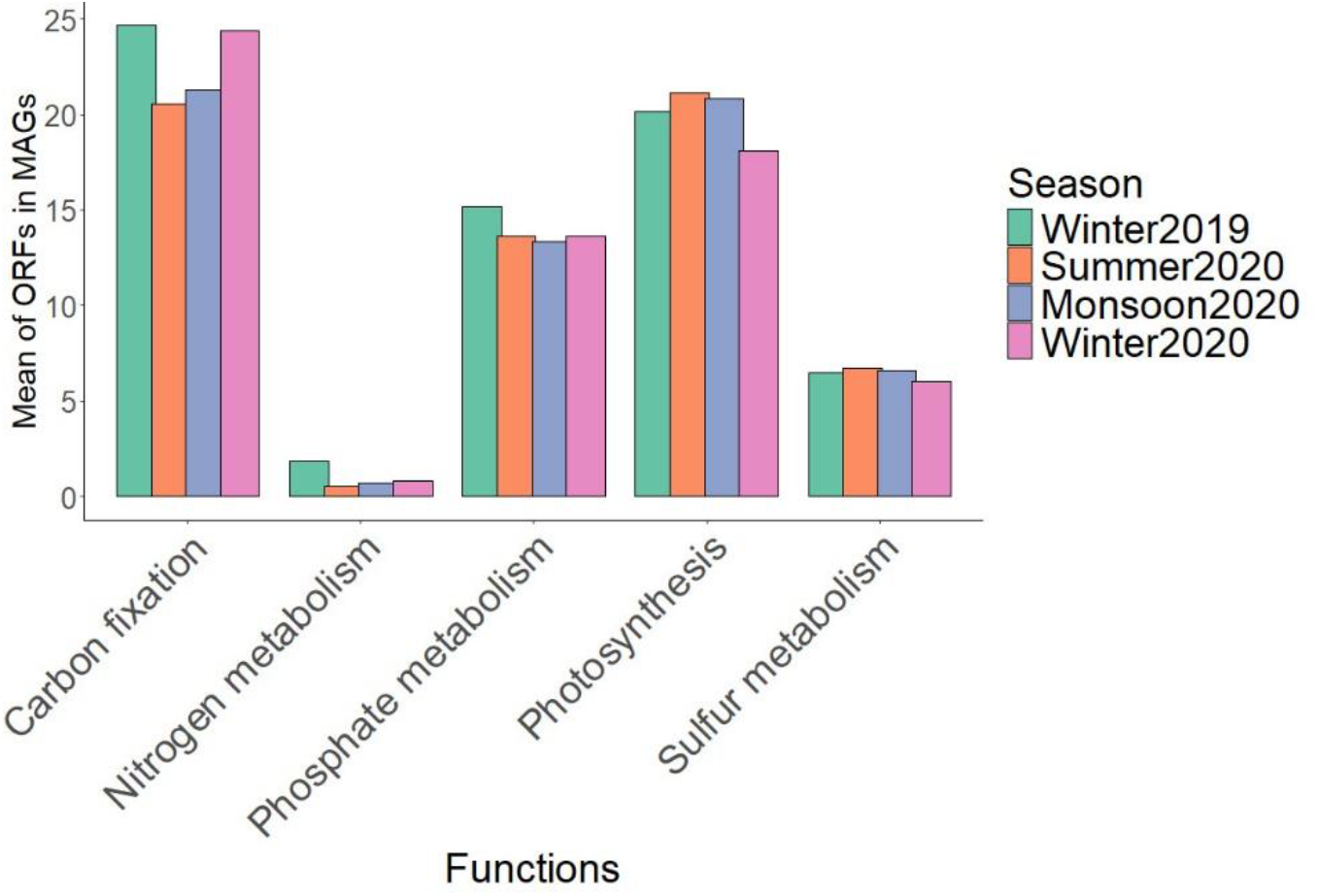
Comparison of the mean number of ORFs for biogeochemical functions in cyanobacterial MAGs

**Figure 6:**
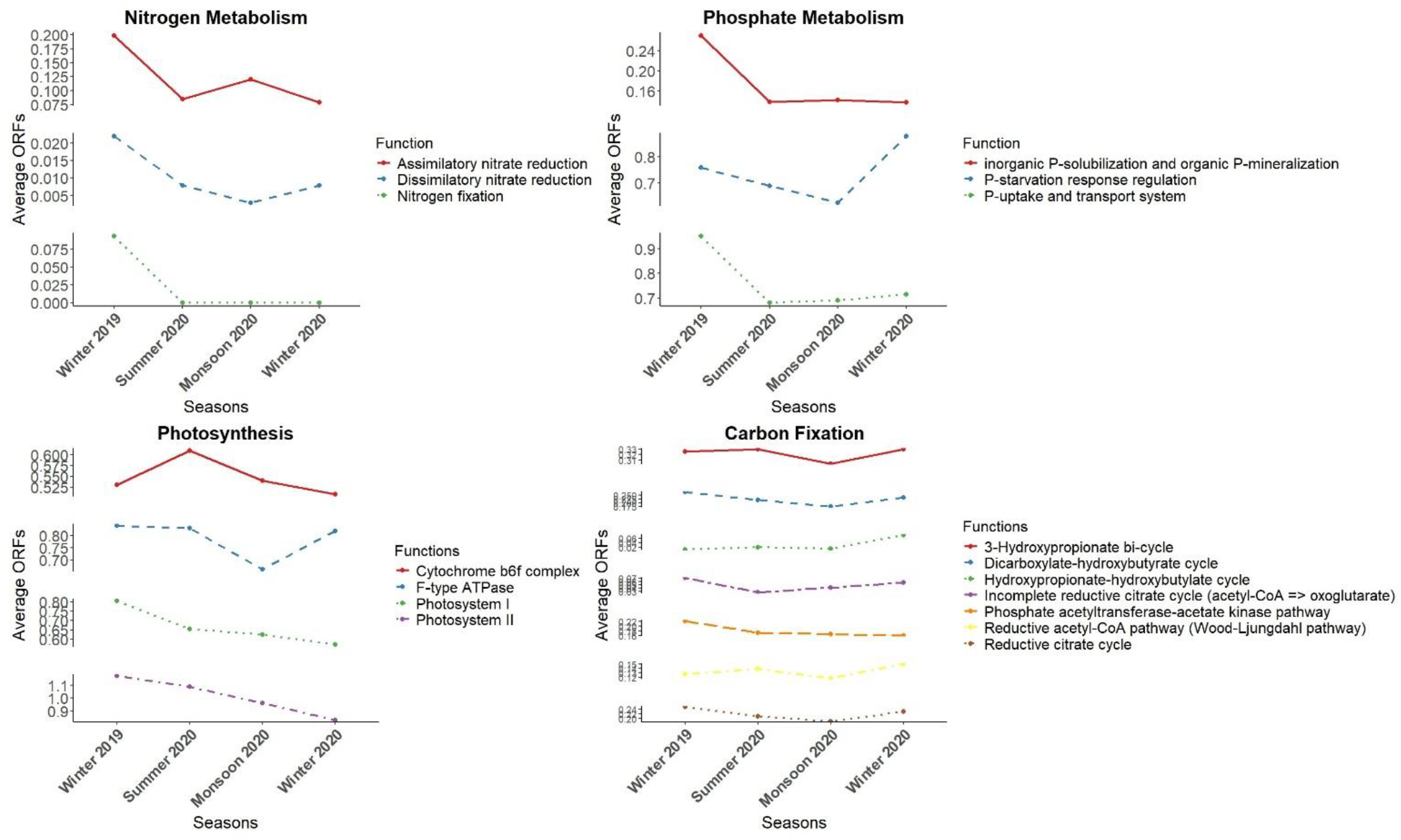
Seasonal variation in average ORFs of cyanobacterial MAGs for six biogeochemically important functions

The plot grouped phosphate metabolism pathways/genes into three categories: phosphate uptake and transport system, inorganic phosphate solubilization, organic phosphate mineralization and phosphate starvation response regulation (Figure 6). The mean ORFs from cyanobacterial MAGs for the first two categories were highest in the winter of 2019. In contrast, the third category of phosphate starvation response regulation was highest in the winter of 2020.

Significantly varying trends of the photosynthetic pathway genes were observed (Friedman test, p-value = 0.0439). The plot shows a greater mean number of ORFs in the winter of 2019 for Photosystems 1 and 2, whereas cytochrome bf6 ORFs were highest in the summer of 2020 (Figure 6). The mean ORFs for F-type ATPases, the enzyme complexes responsible for ATP synthesis during photosynthesis, are almost the same across the seasons, with a minimum in the monsoon of 2020.

The various carbon-fixation pathways across seasons also varied significantly (Friedman test, p = 0.05). Figure 6 shows the temporal change in mean ORFs of the seven major carbon fixation pathways: the reverse tricarboxylic acid (rTCA) cycle/reductive citrate cycle, the incomplete reductive citrate cycle, the 3-hydroxypropionate (3HP) cycle, the dicarboxylate/4-hydroxybutyrate (DC/4HB) cycle, the reductive acetyl-CoA pathway (also known as the Wood‒ Ljungdahl pathway), the 4-hydroxybutyrate/3-hydroxypropionate (4HB/3HP) cycle and the phosphate acetyltransferase‒ acetate kinase pathway. There were 5 MAGs (Table 1) which showed the complete phosphate acetyltransferase-acetate kinase pathway a crucial component of the Wood–Ljungdahl pathway and converts acetyl-CoA to generate ATP. The plot in Figure 6 shows that the season with the highest mean ORFs for different pathways varies. For example, for the four pathways of the reductive citrate cycle, incomplete reductive citrate cycle, phosphate acetyltransferase-acetate kinase pathway and dicarboxylate/4-hydroxybutyrate cycles, the winter of 2019 presented the highest mean ORFs in cyanobacteria. Moreover, the 4-hydroxybutyrate/3-hydroxypropionate and reductive acetyl-CoA pathway had the highest mean ORF in the winter of 2020. The 3-hydroxypropionate (3HP) bicycle has mostly similar mean ORFs across all seasons.

**Table 1:** MAGs showing complete phosphate acetyltransferase-acetate kinase pathway (acetyl-CoA => acetate)

| MAG Id | NCBI accession number | NCBI taxonomy | Score K13788 (pta) | Score K00925 (ackA) | e-value K13788 (pta) | e-value K00925 (ackA) |
| --- | --- | --- | --- | --- | --- | --- |
| 1426476 | JAYKHD000000000 | o__Synechococcales;<br>f__Synechococcaceae;<br>g__; s__ | 660.9 | 350.5 | 5e<sup>-199</sup> | 3.3e<sup>-105</sup> |
| 27796 | JAYKHU000000000 | o__Synechococcales;<br>f__Merismopediaceae;<br>g__Synechocystis; s__ | 939.1 | 571.9 | 3.6e<sup>-283</sup> | 3.3e<sup>-172</sup> |
| 280001 | JAYKHV000000000 | o__Oscillatoriales;<br>f__Oscillatoriaceae;<br>g__Lyngbya; s__ | 1026.1 | 574.6 | 0 | 1.3e<sup>-172</sup> |
| 487179 | JAYKIP000000000 | o__Cyanobacteriales;<br>f__Microcystaceae_A;<br>g__; s__ | 909.3 | 572.3 | 3.8e<sup>-274</sup> | 2.7e<sup>-172</sup> |
| 145596 | JAYKHF000000000 | o__Pseudanabaenales;<br>f__Leptolyngbyaceae;<br>g__Leptolyngbya; s__ | 963.8 | 560.3 | 1.5e<sup>-290</sup> | 1.3e<sup>-168</sup> |

### 3.4 Temporal shifts of biosynthetic gene clusters for secondary metabolite production in the cyanobacterial MAGs

The taxonomic classification of most BGCs revealed that they had both bacterial and eukaryotic origins. The highest proportion of BGCs were taxonomically annotated to belong to the phylum Actinobacteria. The next highest proportion was annotated as Proteobacteria, followed by unassigned phyla Interestingly, a large proportion of BGCs were annotated to be of fungal origin rather than of bacterial origin. This may be due to the undocumented taxonomic classification of many cyanobacterial BGCs or the horizontal transfer of these BGCs.

The BGCs were classified into a maximum of 7 classes. The distribution of the proportion of each BGC class varied from one season to another (chi-square test, p=0.019). In the winters of 2019 and 2020, a total of 7 and 6 BGCs, respectively, were identified, whereas in the summer and monsoon seasons, 5 BGC classes were detected (Figure 7). Terpenes accounted for the greatest proportion among all the secondary metabolites irrespective of season. In the winter of 2019, summer and monsoon seasons, RiPPS (ribosomally synthesized and posttranslationally modified peptides) accounted for the second highest proportion, whereas it was PKS (polyketide synthetases) in the winter of 2020.

**Figure 7:**
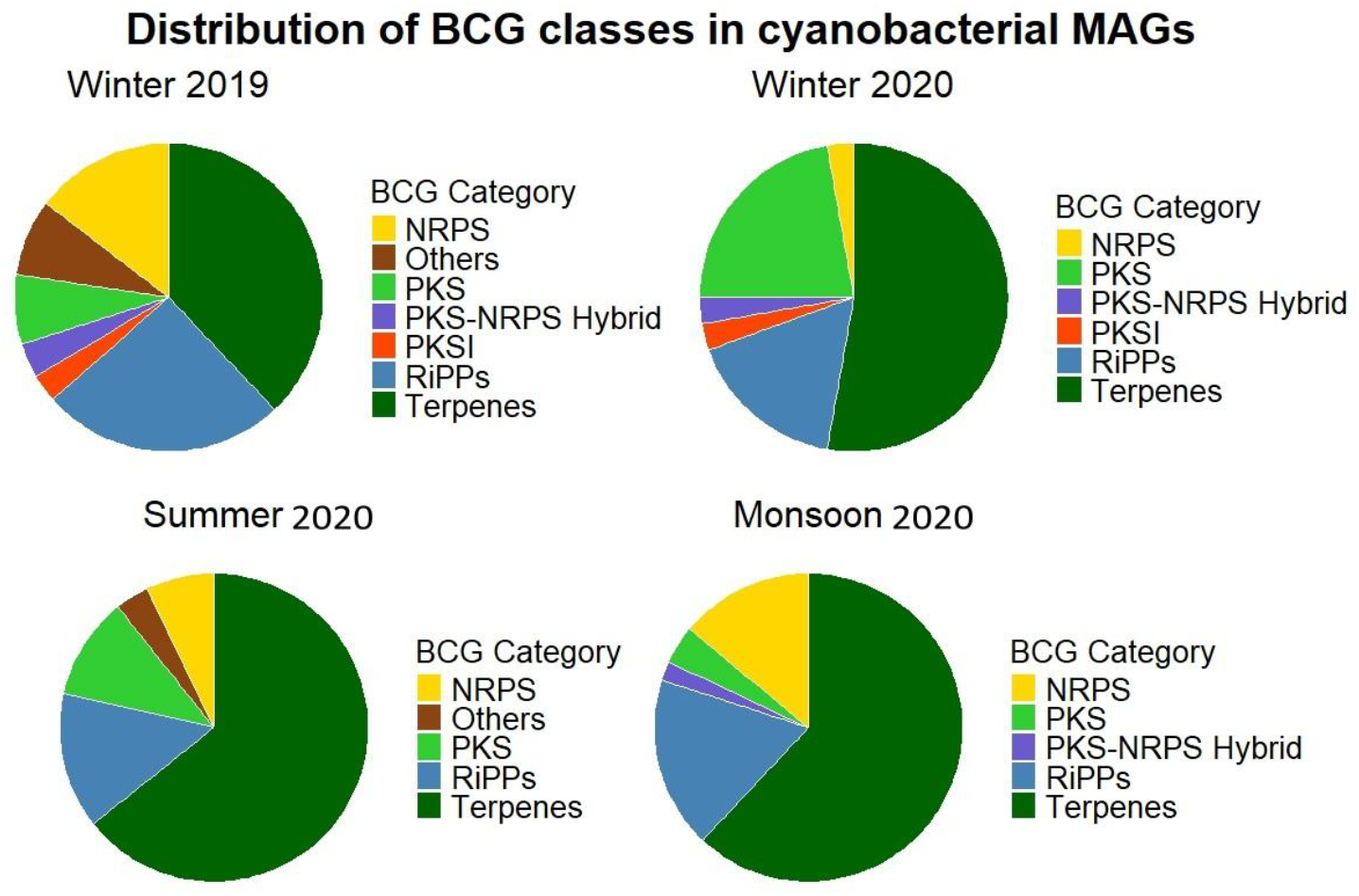
Taxonomic annotation at the kingdom and phylum levels for the biosynthesis-related gene clusters in the cyanobacterial MAGs and temporal distribution of BGC classes in cyanobacterial MAGs

## 4. Discussion

We assembled 83 cyanobacterial MAGs which encoded 11,822 orthogroups. 757 orthogroups of these were species-specific. These MAGs and their orthogroups were present in varying abundances across space and time. Changes in the physiochemical parameters of aquatic ecosystems favour species that can best survive and proliferate under those conditions. Species performance, distribution, and community composition depend on the genomic potential of species to withstand different conditions

### 4.1 Temporal distribution of cyanobacterial MAGs across Chilika Lagoon

We found that certain species were present in all seasons (Cyanobium), but others were season specific. Most interesting finding was Winter of 2019, marked by N:P ratio of less than 16, had 27 unique cyanobacterial MAGs. These MAGs belonged to atleast 11 families. Some of the genera like Snowella (Bukowska et al., 2017), Raphidiopsis (Baxter et al., 2022), Sphaerospermopsis (Kim et al., 2020) and Lyngbya (Osborne et al., 2001) are known to protein toxins in eutrophic conditions (Bukowska et al., 2017). This seasonal specificity of certain cyanobacterial species may be to the optimal environmental conditions for their growth. This further signifies that the preferred cyanobacterial species would have genomic traits to perform crucial biogeochemical functions and produce secondary metabolites (cyanotoxins), which gives them an edge over other cyanobacterial species under the given environmental conditions.

The impact of each abiotic variable on species spatiotemporal dynamics varies (Rubin and Leff, 2007). In our study, salinity emerged as the most significant driver of cyanobacterial species diversity in Chilika Lagoon. Furthermore, salinity also best explains the distribution of COG classes of cyanobacteria. Previous studies on Chilika Lagoon also suggests that salinity is the main factor driving bacterial species composition (Mohapatra et al., 2023). These findings indicate that salinity is a key factor determining which functions encoded by cyanobacterial species are favoured.

In our study, we also observed that patterns of taxonomic or phylogenetic relatedness did not affect the distribution of cyanobacterial MAGs across seasons. This might be because taxonomic relatedness can be decoupled from functional traits due to adaptive evolution and horizontal gene transfer events in microorganisms (Yang, 2021).

### 4.2 Seasonal shifts in major biogeochemical cycling genes of cyanobacteria

Orthogroups are sets of genes that descend from a single gene in the last common ancestor of different species and retain biological function(Emms and Kelly, 2019). We used orthogroups as proxies for functional traits in cyanobacteria. We found that the number of shared orthogroups between pairs of cyanobacterial MAGs was dependent on their taxonomic relatedness.

Chilika Lagoon shows significant heterogeneity in terms of salinity in different sectors: the northern sector (salinity <5), central sector (salinity 6–15), southern sector (salinity >15) and an outer channel connecting to the Bay of Bengal (salinity >30) (Behera et al., 2017). The genomic traits of biogeochemical cycling genes for particular pathways showed temporal variations. Interestingly, only the winter of 2019 was characterized by an N:P ratio of less than 16, which is considered favourable for cyanobacterial growth (Levich, 1996). This season presented more encoded traits for most biogeochemical functions in terms of mean ORFs. This can be correlated with a greater diversity of cyanobacterial species due to optimal growth conditions.

Low biological oxygen and high dissolved oxygen (DO) levels are generally indicative of good water quality in aquatic ecosystems (Kannel et al., 2007) and hence enhanced aerobic reactions by bacteria. Together, these factors might have resulted in the predominance of important functions in the winter of 2019. Additionally, the slightly greater mean ORFs in the winter of 2020 than in the summer may be due to the functions of these ORFs in photosynthesis; hence, examining the distribution of different ORFs might be more informative.

In nitrogen metabolism pathways, DNRA and nitrogen fixation were highest during the winter of 2019, likely due to the nitrogen-to-phosphorus (N:P) ratio being less than 16. The N:P ratio is a critical determinant of nutrient availability, strongly influencing the growth and metabolism of cyanobacteria (Kolzau et al., 2018) and the expression of nitrogen metabolism genes (Kramer et al., 2022). When the N:P ratio falls below 16, it indicates a relative nitrogen deficiency compared to phosphorus. Under such conditions, cyanobacteria experience nitrogen limitation and respond by upregulating genes related to nitrogen acquisition and assimilation. Additionally, cyanobacteria regulate gene expression in nitrogen storage and recycling processes, including the induction of nitrogenase, the enzyme responsible for nitrogen fixation. This enables cyanobacteria to acquire atmospheric nitrogen when dissolved nitrogen levels are insufficient to support growth (Burford et al., 2006). However, nitrogen fixation is associated with slower growth rates compared to periods when both nitrogen and phosphorus are abundant (Kolzau et al., 2018). Consequently, cyanobacteria modulate gene expression to optimize nitrogen utilization under nutrient-limited conditions. In contrast, the monsoon season was characterized by an average N:P ratio of 64, which led to the second-highest mean ORFs for nitrate assimilation and the lowest DNRA. In this scenario, cyanobacteria likely prioritized assimilatory nitrate reduction, as nitrogen was no longer the limiting nutrient (Ohashi et al., 2011). Assimilatory nitrate reduction facilitates the incorporation of available nitrate into organic molecules essential for growth and biosynthesis, including amino acids, proteins, nucleic acids, and other cellular components. Although DNRA plays a role in conserving nitrogen by converting nitrate to ammonium, a bioavailable form for future use (Pandey et al., 2020), its role may be less prominent when nitrogen is abundant in the environment. This explains the lower DNRA observed during the monsoon season, as evident from the comparison plot.

Furthermore, in the winter of 2019, there were greater mean ORFs for phosphate uptake and transportation, inorganic phosphate solubilization and organic phosphate mineralization. The presence of ample phosphate in the environment leads to the uptake of phosphate and its solubilization and mineralization which is beneficial for the metabolic and growth needs of cyanobacteria (Correll, 1999). In contrast, the phosphate starvation ORFs were lower in winter 2019 than in winter 2020, which can be explained by a higher N:P ratio in winter 2020 (17.5) than in 2019 (12.3), implying higher phosphate concentrations in winter 2019 than in winter 2020 compared with the nitrogen concentration in the other seasons. This might have occurred because cyanobacteria uptakes excess phosphate from the environment to cope up with fluctuating availability of phosphate (Solovchenko et al., 2020). Our results for phosphate metabolism genes in cyanobacterial species present in Monsoon 2020 were not as expected. This season was characterized by a high N:P ratio (63.8), which implied a lower phosphate concentration than its nitrogen counterpart and hence increased ORFs for phosphate starvation genes. However, this was not the case in our results, which might be due to other abiotic or biotic interactions that play complex roles in such processes and might be not decipherable from our data.

Even in the photosynthesis pathway, overall, the mean number of ORFs was greater in the winter of 2019. Upon segregating the individual functions of this pathway, and we found that the mean ORF for the cytochrome bf6 complex was greater in the summer. During summer, the light intensity is relatively high and hence affects the rate of photosynthesis. Therefore, increased cytochrome bf6 during summer is expected. Cytochrome bf6 helps mediate the flow of electrons between photosystem I and photosystem II, generating a proton gradient crucial for photosynthesis (Hasan et al., 2013).

The Calvin–Benson–Bassham (CBB) cycle is the primary and only carbon fixation pathway in cyanobacteria, on the basis of the current literature. There are no reports attributing other carbon fixation pathways in cyanobacteria. However, we found that five cyanobacterial MAGs had a complete phosphate acetyltransferase-acetate kinase pathway converting acetyl-CoA to acetate. This is a vital energy conservation step in the Wood–Ljungdahl pathway (Smith et al., 2019). Although none of them had a complete–Ljungdahl pathway, this highlights the possibility of alternate carbon fixation pathways unknown in cyanobacteria.

### 4.3 Seasonal shifts of the biosynthetic gene clusters in cyanobacteria

The secondary metabolites produced by bacteria in their stationary phase (Navarro Llorens et al., 2010) are compounds that do not directly help them in their growth and proliferation. They are generally produced during limiting nutrient conditions and environmental stress and confer ecological functions, such as defense mechanisms, toxin production, signalling, or competitive interactions (Carmichael, 1992) (Maplestone et al., 1992). Cyanotoxins are either non-ribosomal peptide synthetases (NRPSs) or PKSs in nature (Calteau et al., 2014). The cyanobacterial MAGs with a relatively high proportion of PKS or NRPS varied temporally, suggesting a shift in the dominant cyanobacterial species. In the winters of 2019 and during the monsoon season, the NRPS accounted for a greater proportion than did the PKS, whereas the opposite occurred in the winters of 2020 and summer. Furthermore, there were NRPS-PKS hybrids in all seasons except summer. Hence, temporal variation influenced the abundance of cyanobacterial MAGs for a particular class of BGCs. As mentioned earlier, the winter of 2019 was characterized by a lower N:P ratio favourable for cyanobacterial proliferation and blooming. We also found that this season presented a lower proportion of terpenes and a greater proportion of NRPS, PKS, PKSI and their hybrids than the other seasons did. This is because of the presence of cyanobacterial MAGs with the potential for bloom formation. This can be attributed to the role of these BGCs in cyanobacteria and the environmental conditions in which each class is favoured.

## 5. Conclusion

We found that salinity was one of the key abiotic factors that governed the distribution of cyanobacterial species and the functional traits they encoded. The functional traits in terms of pathways for the biosynthesis of secondary metabolites and the biogeochemical cycling of nitrogen, sulfur, phosphate, and carbon and photosynthesis showed temporal variations with changes in the physiochemical parameters of Chilika Lagoon. The kind of cyanobacterial species present in an ecosystem at a given time depends on both abiotic and biotic interactions and is complex and dynamic. However, the overall preference of certain cyanobacterial species to thrive and increase under optimal or environmentally stressed conditions can be deciphered with such studies. Meta transcriptomic data can help provide real-time snapshots of these crucial genes involved in ecosystem function. This further translates to a better comprehension of ecosystem health and the impact of environmental perturbations in the wake of climate change on major primary producers of aquatic ecosystems, Cyanobacteria.

## CRediT authorship contribution statement

Manisha Ray: Conceptualization, Investigation; Methodology, Data curation and analysis, Writing-Original draft preparation, Writing-Reviewing and Editing. Govindhaswamy Umapathy: Conceptualization; Resources; Funding, Supervision, Writing-Reviewing and Editing.

## Declaration of competing interests

The authors declare that they have no competing interests.

## Availability of data and materials

The raw reads analysed during the current study are available in the NCBI SRA under the BioProject accession number PRJNA691704.

## Acknowledgements

We are grateful to Dr. Meghna Krishnadas for her valuable comments and guidance on the statistical analysis and manuscript review. We are also thankful to the Chilika Development Authority, Balugaon, Odisha, for offering logistical support for sample collection. Additionally, we appreciate Dr. Shivakumara Manu’s assistance with the metagenomic assemblies of cyanobacteria.

## Funding

We acknowledge the support provided to G.U. from the Department of Biotechnology (DBT), Govt. of India through Grant No. BT/PR29032/FCB/125/4/2018 for this study, which also funded a partial PhD fellowship for M.R.

## Supplementary information

Supplementary 1: Plot showing the number of orthogroups in each cyanobacterial MAG

Supplementary 2: Heatmap depicting the number of orthogroups shared between pairs of cyanobacterial MAGs

Supplementary 3: Plot showing the Pearson correlation between the abiotic variables of Chilika Lagoon

